# MicroRNAs Cause Accelerated Decay of Short-Tailed Target mRNAs

**DOI:** 10.1101/763367

**Authors:** Timothy J. Eisen, Stephen W. Eichhorn, Alexander O. Subtelny, David P. Bartel

## Abstract

MicroRNAs (miRNAs) specify recruitment of deadenylases to mRNA targets. Despite this recruitment, we find that miRNAs have almost no effect on steady-state poly(A)-tail lengths of their targets in mouse fibroblasts, which motivates acquisition of pre-steady-state measurements of the effects of miRNAs on tail lengths, mRNA levels, and translational efficiencies. Effects on translational efficiency are minimal compared to effects on mRNA levels—even for newly transcribed target mRNAs. Effects on target mRNA levels accumulate as the mRNA population approaches steady state, whereas effects on tail lengths peak for recently transcribed target mRNAs and then subside. Computational modeling of this phenomenon reveals that miRNAs cause not only accelerated deadenylation of their targets but also accelerated decay of short-tailed target molecules. This unanticipated effect of miRNAs largely prevents short-tailed target mRNAs from accumulating despite accelerated target deadenylation. The net result is a nearly imperceptible change to the steady-state tail-length distribution of targeted mRNAs.

**Highlights:**

- miRNAs cause accelerated decay of short-tailed target molecules
- This accelerated decay largely prevents accumulation of short-tailed target mRNAs
- miRNAs are similarly effective on short-lived and long-lived target mRNAs
- In 3T3 cells, miRNA effects on translation are negligible—even for nascent mRNA

## Introduction

The miRNA pathway targets mRNAs from most human genes (Friedman et al., 2009). Each miRNA associates with an Argonaute (AGO) protein to form a silencing complex. Within this complex, the miRNA pairs to regulatory sites within mRNAs (Bartel, 2018), whereas AGO interacts with TNRC6/GW182, which in turn recruits the PAN2–PAN3 and the CCR4–NOT deadenylase complexes, with the CCR4–NOT complex being more consequential (Chen et al., 2009; Braun et al., 2011; Chekulaeva et al., 2011; Fabian et al., 2011; Christie et al., 2013; Huntzinger et al., 2013). This recruitment explains the observation that miRNAs accelerate deadenylation of reporter mRNAs (Giraldez et al., 2006; Wu et al., 2006; Chen et al., 2009) and is thought to underlie the widespread influence of miRNAs on post-transcriptional gene expression (Jonas and Izaurralde, 2015; Bartel, 2018).

The regulatory effects of miRNAs can differ, depending on the consequences of tail shortening in the posttranscriptional regulatory regime operating in the cell (Subtelny et al., 2014). For example, in early zebrafish embryos, where poly(A)-tail length influences translational efficiency but not mRNA stability, miRNAs cause reduced translational efficiency with little effect on mRNA stability (Bazzini et al., 2012; Subtelny et al., 2014). However, in gastrulating embryos, which are undergoing a developmental shift in the nature of translational control, such that poly(A)-tail length no longer influences translational efficiency but instead influences mRNA stability, miRNAs predominantly cause reduced mRNA stability (Bazzini et al., 2012; Subtelny et al., 2014). Likewise, in the post-embryonic systems that have been studied to date, mRNA destabilization is the principal mode of miRNA-mediated repression of endogenous messages, with a minor contribution from translational repression sometimes also detected (Baek et al., 2008; Hendrickson et al., 2009; Guo et al., 2010; Eichhorn et al., 2014). This minor contribution from translational repression is not attributable to poly(A)-tail shortening, as poly(A)-tail length is not coupled with translational efficiency in these post-embryonic systems (Subtelny et al., 2014).

The lack of substantial effects of miRNAs on translational efficiency observed in post-embryonic systems at steady state does not rule out the idea that miRNAs might have a greater effect on translation soon after they encounter their targets. Investigations of this idea involving the dynamics of miRNA-mediated repression have yielded different results, depending upon the experimental approach. One approach has been to rapidly induce the expression of a reporter mRNA and measure its initial repression by a highly expressed constitutive miRNA (Bethune et al., 2012; Djuranovic et al., 2012), and the other approach has been to rapidly induce the expression of a miRNA and measure the repression of cellular mRNAs (Eichhorn et al., 2014). Translational repression seems to be much more prominent at early time points of the first approach compared to those of the second. One explanation for this discrepancy is that repression can be more readily detected at early time points if the miRNA is already at full strength at these time points. A second explanation is that reporter mRNAs might respond differently than endogenous cellular mRNAs. Analyses of the initial repression of newly transcribed endogenous mRNAs could distinguish between these possibilities.

Pre-steady-state analyses of miRNA effects might also provide mechanistic insight into miRNA-mediated mRNA destabilization. For instance, if the miRNA pathway acts independently of other decay pathways, then it would be expected to contribute less to the decay rate of rapidly decaying targets and, by extension, have greater influence on the expression level of long-lived mRNAs. Alternatively, if the miRNA pathway acts together with and reinforces other mRNA-decay pathways, then it would be expected to act more rapidly on mRNAs that are already decaying more rapidly, and contribute more equally to the destabilization of mRNAs regardless of their basal decay rate. Analyses of steady-state data seem to indicate that mRNAs with shorter half-lives are less susceptible to miRNA-mediated destabilization, which suggests at least some independence (Larsson et al., 2010). However, steady-state data cannot speak to the question of whether the miRNA pathway is acting more rapidly on some mRNAs than on others, and the results leave open the possibility of concerted action with other pathways.

We have developed an experimental and computational framework to study the dynamics of deadenylation and decay of endogenous mRNAs of thousands of genes (Eisen et al., 2019). A conclusion of that study, performed in parallel with the current study, is that once deadenylation has proceeded to the point that mRNAs have short tails, these short-tailed mRNAs degrade at dramatically different rates. Moreover, mRNAs that have undergone more rapid deadenylation decay more rapidly upon reaching short tail lengths (Eisen et al., 2019). These results bring up the possibility that miRNA-mediated mRNA destabilization might have a second mechanistic component: In addition to the well-established role of promoting deadenylation, miRNAs might also cause more rapid decay of short-tailed target mRNAs. To investigate this possibility and to explore other questions that have awaited a pre-steady-state analysis of the global effects of miRNAs on endogenous mRNAs of the cell, we extended the experimental and computational framework of our concurrent study to also query translational control. With this extended framework, we set out to investigate the effects of miRNAs on the dynamics of cytoplasmic mRNA metabolism, simultaneously examining their effects on tail lengths, translation, and decay of target mRNAs.

## Results

### Very Slight Effects of miRNAs on Bulk mRNA Poly(A)-Tail Lengths

Because miRNAs specify recruitment of deadenylation machinery to their mRNA targets (Jonas and Izaurralde, 2015), we first examined the global effects of miRNAs on the steady-state tail lengths of mammalian mRNAs, making use of an improved version of poly(A)-tail length profiling by sequencing (PAL-seq, (Eisen et al., 2019)). In this experiment, miR-1 was induced for 48 h in a clonal 3T3 cell line previously engineered to enable inducible miR-1 expression (Eichhorn et al., 2014), and tail lengths were profiled in these cells expressing miR-1 as well as cells grown in parallel without miR-1 induction. Despite robust reduction of both mRNA and ribosome-protected fragments (RPFs) of predicted miR-1 targets, as measured using RNA-seq and ribosome-footprint profiling (Ingolia et al., 2009), respectively, the mean tail-length change for these targets was < 0.1 nt, which was not statistically significant (Figure 1A left; p *=* 0.83). When focusing the analysis on a set of top-repressed targets, defined as those repressed by at least 25% in a prior experiment in which miR-1 was induced (Eichhorn et al., 2014), mean tail-length change was still < 0.1 nt (Figure 1A right; p = 0.90). Essentially no change was also observed for predicted miR-155 targets after inducing miR-155 (Figure 1B; < 1 nt change for both predicted targets and top targets, p = 0.35 and 0.46, respectively).

**Figure 1.**
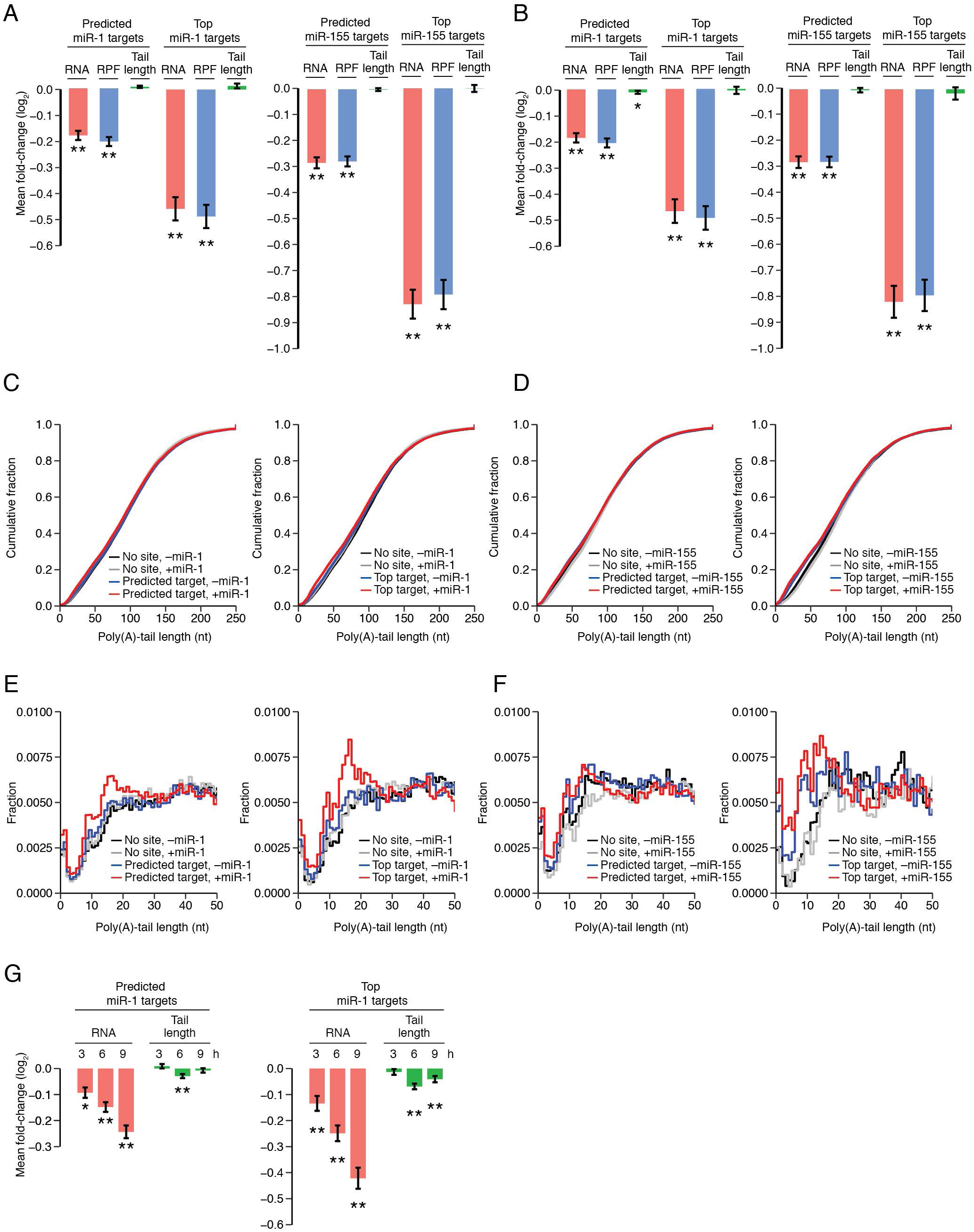
Very Small Effect of miRNAs on Steady-State Poly(A)-Tail Lengths. (A) Impact of miR-1 and miR-155 (left and right, respectively) on RNA abundance, RPF abundance, and mean poly(A)-tail length of their predicted targets (n = 657 and 466 for miR-1 and miR-155, respectively) and top predicted targets (n = 133 and 65, respectively). Values were normalized to those of mRNAs with no site to the induced miRNA (n = 2170 and 3501, respectively). Significant changes are indicated with asterisks below the bar (* p ≤ 0.05; ** p ≤ 0.001, one-tailed t-test). Error bars are the standard error of the mean (SEM), propagating the error for the SEM of the no-site mRNAs. (B) Impact of miR-1 and miR-155 on RNA abundance, RPF abundance, and mean poly(A)-tail length of their predicted targets (n = 643 and 417 for miR-1 and miR-155, respectively) and top predicted targets (n = 130 and 57, respectively) when using a protocol that does not deplete very short or highly modified tails. For each gene, PAL-seq data were used to determine the length distribution of tails ≥ 50 nt, and data generated using a protocol without a splinted ligation were used to determine both the length distribution of tails < 50 nt and the fraction of tails < 50 nt. Values were normalized to those of mRNAs with no site to miR-1 or miR-155 (n = 2131 and n = 3138, respectively); otherwise, as in (A). (C) Very small influence of miR-1 on the steady-state tail-length distribution of its predicted targets. Left: cumulative distributions of poly(A)-tail lengths in miR-1– uninduced and miR-1–induced cells (–miR-1 and +miR-1, respectively) for predicted miR-1 targets (n = 643) and a set of no-site mRNAs with the same distribution of 3′ UTR lengths (n = 626). To ensure that the tail-length distributions were not unduly influenced by a few highly-expressed mRNAs, values for mRNAs with more than the median number of poly(A) tags were down-weighted such that they contributed to the distribution as much as values from the mRNA with the median value. Right: As on the left, but plotting the results for top predicted targets of miR-1 (n = 130) and a set of no-site mRNAs matched for UTR length (n = 130). Results are from datasets used in (B), which avoided depletion of mRNAs with very short or highly modified tails. (D) Very small influence of miR-155 on the steady-state tail-length distributions of its predicted targets. Analysis is as in (C) but showing the effect of inducing miR-155 on tail lengths of its predicted targets (n = 417 and 57 for predicted and top targets, respectively) and matched controls (n = 417 and 57, respectively). (E) Influence of miR-1 on the steady-state tail-length distribution of short-tailed mRNAs. Analysis as in (C), except showing results for predicted targets (left) and top predicted targets (right) with tails ≤ 50 nt as a histogram. (F) Influence of miR-155 on the steady-state tail-length distributions of short-tailed mRNAs. Otherwise as in (E). (G) Reanalysis of data monitoring mRNA levels and tail lengths after transfecting miR-1 into HeLa cells (Chang et al., 2014). Changes in mRNA abundance of predicted miR-1 targets (n = 354) and top predicted miR-1 targets (n = 140) normalized to no-site controls (n = 908). For each time point, changes were normalized by the corresponding median log_2_ fold-change of the 0 h time point. Otherwise as in (A).

These analyses used datasets generated from a splint ligation to the poly(A) tail, which was expected to be less efficient for mRNAs with tail lengths < 8 nt, and although the protocol was able to detect mRNAs with tails modified with a single terminal U, albeit at reduced efficiency, it was not designed to detect mRNAs with tails with an oligo(U) or other modification (Eisen et al., 2019). Reasoning that this method might obscure miRNA-induced tail-length changes for mRNAs with very short or extensively modified tails, we performed the same experiments, except the splint ligation was replaced with single-stranded ligation. With these additional data, mean tail length changes were all < 1 nt, and statistically significant for only one of the four comparisons (Figure 1B, p = 0.011 for predicted targets of miR-1).

Having found that inducing expression of a miRNA caused essentially no detectable differences in the mean tail lengths of its predicted targets, we compared single-molecule tail-length distributions to query whether miRNAs altered these distributions in ways that did not affect mean values. Such differences would be expected if miRNAs preferentially direct the deadenylation of shorter tails, as might be inferred from reports that 1) miRNA function depends more on recruitment of the Ccr4–Not deadenylase complex than it does on recruitment of the Pan2–Pan3 deadenylase complex (Behm-Ansmant et al., 2006; Tritschler et al., 2010), and 2) the Ccr4–Not deadenylase complex acts at a later, more processive step of mRNA deadenylation than does the Pan2–Pan3 complex (Yamashita et al., 2005). Contrary to this expectation, induction of either miR-1 or miR-155 had almost no influence on the distribution of single-molecule tail-length measurements for all predicted targets or top-repressed targets of each miRNA (Figure 1C–D).

Analysis of tail-length distributions of mRNAs from different genes showed that for 116 of 1060 predicted targets, induction of the miRNA shortened the tail-length distribution (Kolmogorov–Smirnov (K–S) test, p < 0.01). However, this group of 116 was not much larger than the 81 predicted targets for which induction of the miRNA lengthened the tail-length distribution. Moreover, for 83 of 1041 no-site controls, induction of the miRNA also significantly shortened the tail-length distribution. Taken together, these analyses hinted at a signal, albeit of very low magnitude, for miRNA-dependent shortening of tail lengths. Indeed, closer inspection of the distributions of Figure 1C–D revealed a slight increase in short-tailed isoforms of predicted targets in the presence of the miRNA, which was most pronounced at tail lengths of 7–20 nt (Figure 1E–F)—a range resembling that of the build-up generally observed for short-tailed isoforms of short-lived mRNAs (Eisen et al., 2019). However, for each comparison, these increases involved only a small fraction (0.001–0.021) of the target mRNAs.

### Time-Resolved Measurements of miRNA-Mediated Changes

How then, might this low-magnitude effect observed for endogenous mRNAs be reconciled with both the established role for the poly(A) tail in stabilizing mRNAs (Goldstrohm and Wickens, 2008) and the established role for miRNAs in directing poly(A)-tail shortening of both reporter mRNAs in mammalian cells and endogenous mRNAs in early zebrafish embyros (Giraldez et al., 2006; Wu et al., 2006; Subtelny et al., 2014)? A clue to this riddle comes with the report of tail-length shortening measured 3, 6, and 9 h after miRNA transfection into HeLa cells (Chang et al., 2014). Further analyses of these tail-length changes revealed statistically significant, albeit small, tail-length shortening of predicted targets in this system in which the transcriptome is still adjusting to the recent transfection of a miRNA and has not yet reached a new steady state (Figure 1G, mean changes of 1.9 and 4.0 nt for all predicted targets and top predicted targets, respectively, 6 h after transfection). Perhaps measurements obtained at steady-state are not suitable for detecting miRNA effects on tails, and pre-steady-state measurements must be examined instead.

To understand how the miRNA pathway leverages the relationship between tail lengths and mRNA turnover, we set out to build an experimental and analytical framework for the global study of pre-steady-state miRNA-mediated changes in tail-lengths and mRNA levels. To address lingering questions related to the dynamics of miRNA-mediated translational repression and the potential relationship between these dynamics and those of mRNA decay, we also designed the experiments to detect early miRNA-mediated changes in translational efficiency. Our framework built upon the one that we concurrently developed to study the dynamics of cytoplasmic mRNA metabolism more generally (Eisen et al., 2019). Indeed, the cells used in that study had been engineered to have the ability to inducibly express either miR-1 or miR-155, which allowed data generated in the uninduced condition to be used in both that study and in the current study. In the other study, data generated in the uninduced condition were used to examine the dynamics of mRNA deadenylation and decay, whereas in the current study, those data provided the baseline for comparison to analogous datasets concurrently generated using the same cell lines grown for 48 h in presence of doxycycline, which induced either miR-1 or miR-155, depending on the cell line. In these conditions, mRNAs emerged into a cytoplasm in which the induced miRNA was expressed at a level comparable to those of the highest-expressed endogenous miRNAs (Eichhorn et al., 2014).

Study of pre-steady-state mRNA dynamics, and the influence of miRNAs on these dynamics, required datasets reporting mRNA abundance, poly(A)-tail length, and translation for mRNAs of different ages. To generate these datasets, we performed continuous metabolic-labeling experiments in which the uridine analog 5-ethynyl uridine (5EU) was added to cells, and after different time intervals, cytoplasmic RNA was harvested for analysis of mRNA abundances, translational efficiencies, and tail lengths (Figure 2A). These analyses were each designed to report on only RNA that had been produced after 5EU addition, which was separated from pre-existing RNA by virtue of the alkyne moiety of 5EU; this moiety can be specifically and efficiently biotinylated using click chemistry, thereby enabling isolation of metabolically labeled RNA (Jao and Salic, 2008; Eisen et al., 2019; Kwasnieski et al., 2019). When accounting for the time required for 5EU incorporation, mRNA processing, and mRNA export, most mRNAs isolated from the 40 min time interval had been in the cytoplasm for 0–10 min, whereas mRNAs isolated for longer time intervals had a correspondingly broader range of ages (Eisen et al., 2019). For example, most mRNAs isolated from the 1 h time interval had been in the cytoplasm for 0–30 min. For time-resolved tail-length profiling, 5EU-containing RNA was isolated prior to PAL-seq, whereas for time-resolved ribosome-footprint profiling, 5EU-containing RPFs were isolated and sequenced (Figure 2B). Likewise, to generate RNA-seq data comparable to the RPF data, we fragmented mRNA to the size of RPFs, and isolated and sequenced 5EU-containing mRNA fragments (Figure 2B). Generation of these RPF and RNA-seq datasets required purification of fragments containing a single 5EU, a task greatly facilitated by the high efficiency of 5EU biotinylation (Figure S2).

**Figure 2.**
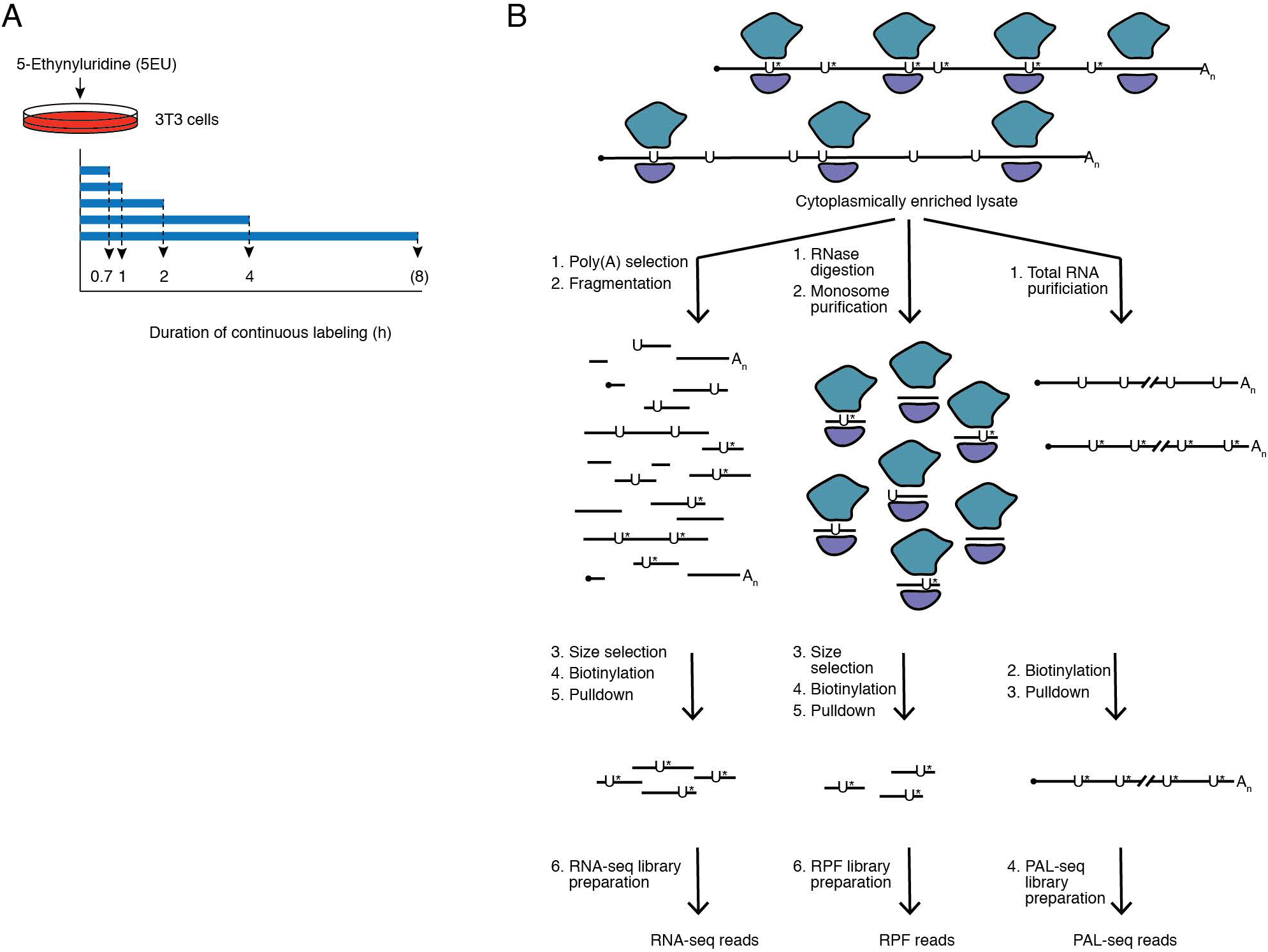
Profiling Dynamics of miRNA Effects. (A) Schematic of 5EU metabolic-labeling. The time point in parentheses indicates that experiments with miR-155–inducible cells but not miR-1–inducible cells included an 8 h labeling period. (B) Schematic of the global assays applied to mRNAs labeled with 5EU (U*). All samples were harvested as cytoplasmically enriched lysates, and for each cell line and condition, all three libraries for each time interval were prepared from the same starting sample.

### Effect of miRNAs on Tail-Lengths but not on Translational Efficiency of Young mRNAs

In cells in which miR-1 was induced, we observed significant decreases in both mRNA and RPFs for predicted miR-1 targets, and the magnitude of these decreases accrued with longer labeling periods (Figure 3A). When considering only the top predicted targets, repression was both more pronounced at each time period and reached statistical significance at earlier time periods (Figure 3A, bottom). Analogous results were observed upon miR-155 induction (Figure 3B). Translational repression, measured as changes in RPFs beyond those observed for mRNA abundance, was difficult to detect in either the miR-1 or the miR-155 experiment. Thus, miRNA-mediated translational repression of cellular mRNAs either does not occur in these cells or is too subtle to be reliably detected, even at short time periods. The observation of robust mRNA destabilization with no sign of translational repression implied that early translational repression is not required to achieve destabilization of endogenous mRNAs.

**Figure 3.**
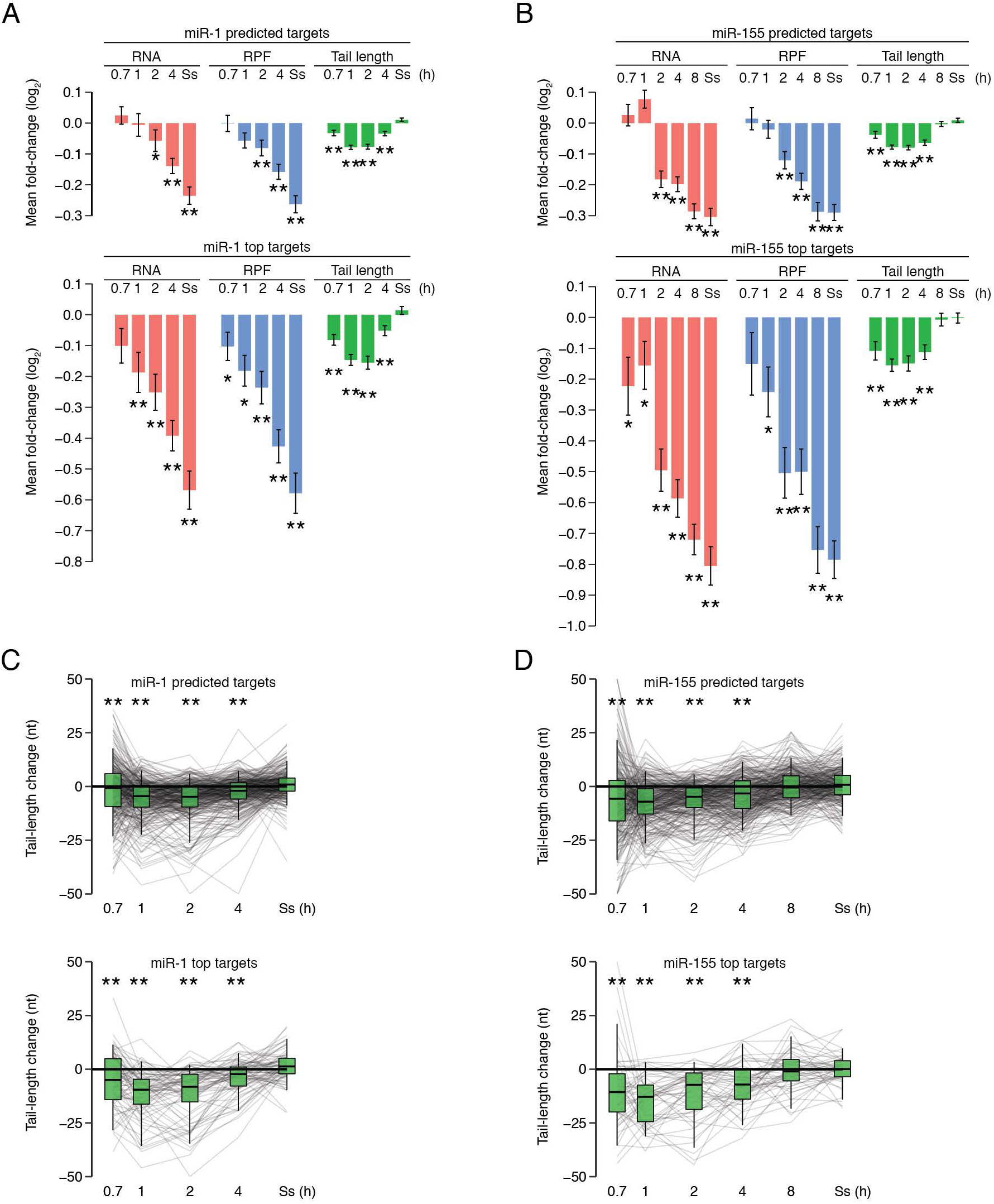
Influence of miRNAs on the Dynamics of mRNA Metabolism. (A) The dynamics of miR-1–mediated repression. Plotted are mean fold-changes in mRNA abundance, RPF abundance, and mean poly(A)-tail length of mRNAs captured after labeling with 5EU for the indicated time intervals, comparing the values observed with and without miR-1 for predicted miR-1 targets (top) and top miR-1 targets (bottom), normalized to those observed for no-site mRNAs (n = 290, 70, and 684, respectively). Results are shown for mRNAs that passed the cutoffs for RNA, RPF, and tail-length measurements at all time intervals. Normalization, error bars, and significance testing were as in Figure 1A. (B) The dynamics of miR-155–mediated repression. Analysis was of values for predicted miR-155 targets (top) and top miR-155 targets (bottom) normalized to those observed for no-site mRNAs (n = 251, 41, and 1470, respectively); otherwise, as in (A). (C) Tail length changes for predicted and top predicted targets of miR-1. Lines connect results for mRNAs from the same gene, and box plots show the spread of the distribution (line, median; box, 25^th^–75^th^ percentiles; whiskers, 5^th^–95^th^ percentiles). Otherwise as in (A). (D) Tail length changes for predicted and top predicted targets of miR-155. Otherwise as in (C).

In contrast to miRNA-mediated translational repression, miRNA-mediated tail shortening was readily detected in the pre-steady-state time periods, and moreover it exhibited very different dynamics from those observed for mRNA and RPF changes. For the predicted targets of miR-1, statistically significant tail shortening was detectable after only 40 min of labeling, peaked between 1 and 2 h, and then abated such that no significant miRNA-mediated tail shortening was observed at steady state (Figure 3A, top). The same behavior was observed for the predicted targets of miR-155, with inclusion of the 8 h labeling interval indicating that tail-length changes abated by this time (Figure 3B, top). Median tail-length changes observed after 1 h of labeling for predicted targets of miR-1 and miR-155, were 4.4 and 7.1 nt, respectively (Figures 3C–D). When considering only the top predicted targets for each miRNA, essentially the same dynamics were observed (Figures 3A–B, bottom), but with larger median changes of 9.6 and 12.8 nt observed after 1 h of labeling for top targets of miR-1 and miR-155, respectively (Figure 3C–D). In both experiments, statistically significant tail-length changes either preceded or occurred concurrently with statistically significant RNA and RPF changes (Figure 3A–D).

### An Analytical Framework for miRNA-mediated mRNA Dynamics

To explain the pattern of tail-length changes observed for miRNA targets, we used a computational model that describes the dynamics of deadenylation and decay for thousands of mRNAs (Eisen et al., 2019). This model, which had been fit to the continuous-labeling data for tail lengths and mRNA abundance in the absence of miRNA induction, was used to simulate the effects of miRNAs over time using three alternative approaches that each tested a different explanation for the fold-change in steady-state abundance for the predicted targets (Figure 4A–B). One explanation tested was the prevailing notion that miRNAs act to destabilize mRNAs by causing more rapid deadenylation of their targets. To test this possibility, we increased the deadenylation rate constants to achieve the fold-changes in steady-state abundance observed with miRNA induction and then used the model with and without these increased values to simulate the miRNA-mediated mRNA and tail-length changes observed in the continuous-labeling experiment, including a 35 min offset to account for nuclear processing and export. Although attributing the entire miRNA effect to increased rates of deadenylation resulted in simulated mRNA-abundance changes that roughly matched the observed changes, the simulated mean tail-length changes diverged from the observed measurements at later time intervals (Figure 4A–B, green). Attributing the entire miRNA effect to only increased rates of short-tailed mRNA decay (called “decapping” in the model) resulted in even greater divergence between the simulated and observed tail-length changes, as more rapid decay of short-tailed mRNAs led to increased rather than decreased mean tail lengths (Figure 4A–B, red). By contrast, when miRNAs were allowed to increase the rates for both deadenylation and short-tailed mRNA decay, the best concordance was achieved; as for the observed mean tail-length changes, simulated changes peaked at short time periods, and then as short-tailed targets decayed more rapidly and a new steady state was established, the effects on mean tail length disappeared (Figure 4A–B, blue).

**Figure 4.**
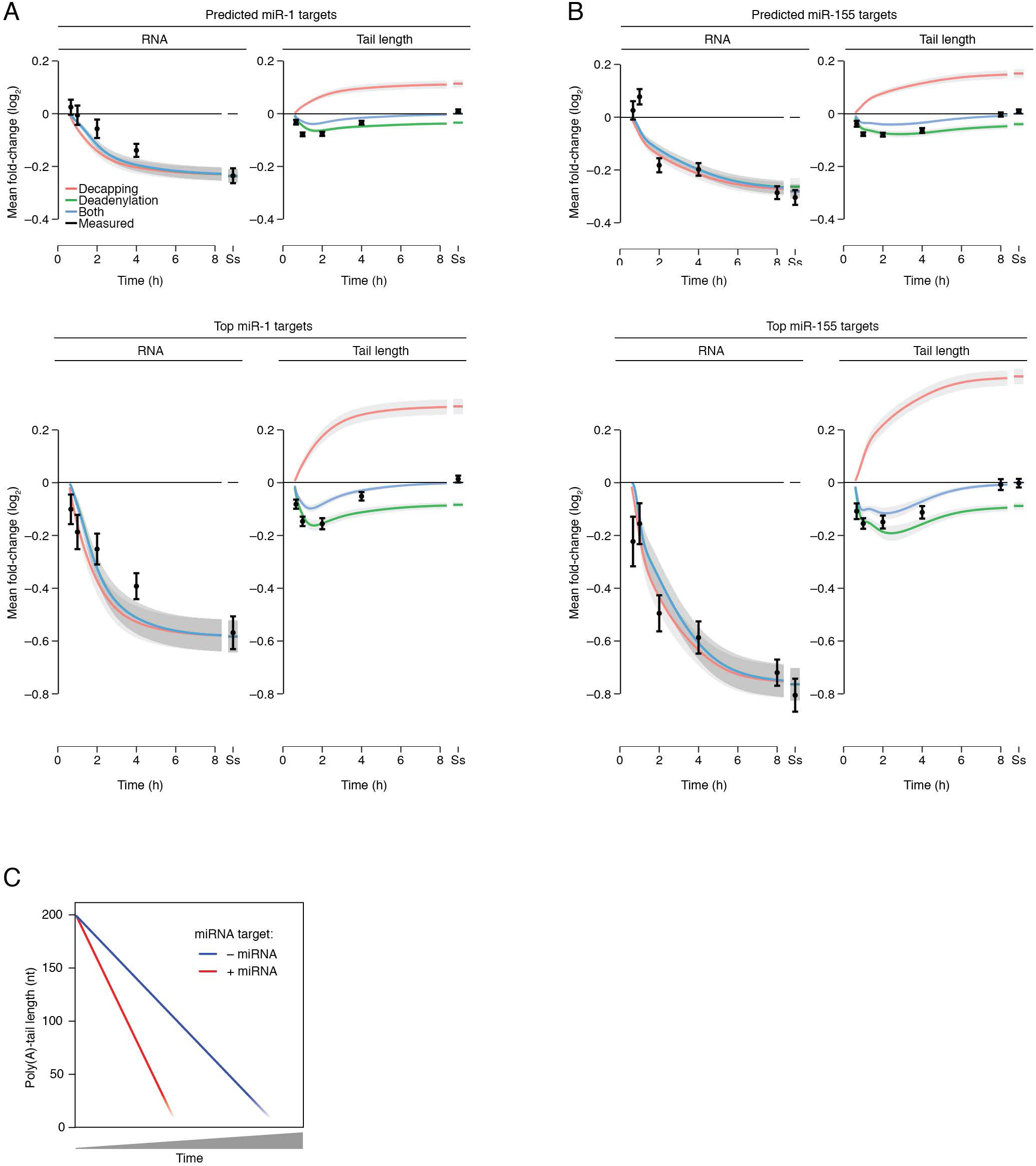
miRNAs Affect Deadenylation and Short-Tailed mRNA Decay. (A) Simulations of effects of miR-1 on its predicted targets (top panels) and top predicted targets (bottom panels). Effects of miRNA were simulated starting with fitted parameters describing initial tail lengths, production rates, deadenylation rates and decapping rates of predicted targets over time without miRNA induction (Eisen et al., 2019). Effects of the miRNA were then modeled as changes in either the decapping rate constant, the deadenylation rate constant, or both rate constants. These changes in rate constants were fit for mRNAs of each gene, minimizing the squared difference between the simulated and measured fold-change in steady-state mRNA abundance. Using these fitted values, the mean fold-changes in mRNA abundance (left) and mean tail lengths (right) were simulated and plotted over time with a 35 min offset to account for nuclear processing and export. For each simulation, the rate constant or combination of rate constants that was changed is indicated (key), with shading showing the SEM. Numbers of predicted targets in each set are as in Figure 3A. (B) Simulations of miRNA effects of miR-155 on its predicted targets (top panels) and top predicted targets (bottom panels). Otherwise as in (A), with numbers of targets as in Figure 3B. (C) Schematic illustrating why miRNAs minimally affect steady-state tail lengths. In the absence of the miRNA, the mRNA exits the nucleus and undergoes deadenylation and then ultimately decay, once its tail becomes short (blue). In the presence of the miRNA, the mRNA target undergoes faster deadenylation, and once its tail is short, faster decay (red). With this more rapid decay of short-tailed isoforms, the targeted mRNA transits through the same distribution of tail lengths as it would have if it were not targeted—but just more rapidly. As a result, the weighted average of the populated tail-length states at steady state (the mean tail length) is unaffected, and the effect of the miRNA on tail length is revealed only when observing pre-steady–state kinetics.

These results indicate that miRNAs cause not only an increase in deadenylation rates of their targets but also an increase in rates at which short-tailed targets are degraded. The increased rate of short-tailed target degradation prevents a steady-state buildup of short-tailed isoforms; without this additional effect of miRNAs, the miRNA-mediated increase in deadenylation rate would lead to a corresponding increase in steady-state short-tailed isoforms, thereby decreasing steady-state mean tail length. With this additional effect of miRNAs, targeted mRNAs can transit through essentially the same distribution of tail lengths, spending the same proportion of time at each tail length in the presence of the miRNA as they would have in its absence (Figure 4C). Thus, this additional effect of miRNAs explains how miRNAs can cause accelerated target deadenylation with minimal influence on tail-length distributions (Figure 1) and why the effect of the miRNA on tail length is revealed only when examining the dynamics of mRNA metabolism.

### Influence of mRNA Half-Life on the Dynamics of miRNA-Mediated Repression

One factor proposed to influence the impact of miRNAs on the expression of their targets is the basal decay rate of the target (Larsson et al., 2010). Indeed, if the induced miRNAs acted independently of the basal decay factors (which include other miRNAs), then the decay rate of a miRNA target would be the sum of the basal decay rate of the target and the decay rate imposed by the induced miRNA, in which case, the targets with faster basal decay rates would experience diminished fold-changes upon miRNA induction compared to targets with slower basal decay rates.

To examine this possibility, we looked at the relationship between steady-state repression of predicted targets and their basal half-lives. Although short-lived predicted targets of miR-1 appeared somewhat less repressed at steady state, the opposite trend was observed for predicted targets of miR-155 (Figure 5A, *R*_s_ = –0.13 and 0.15, respectively, with p = 0.007 and 0.008). This equivalent repression of targets with short and long half-lives could not be explained by more effective target recognition of short-lived mRNAs offsetting less effective target repression of these mRNAs (Figure S3). These results did not support the model in which miRNAs independently add to the basal decay rates of their targets. Instead, they supported a model in which the miRNA silencing complex acts together with the basal decay factors, such that mRNAs with faster basal decay are no less influenced by miRNA targeting.

**Figure 5.**
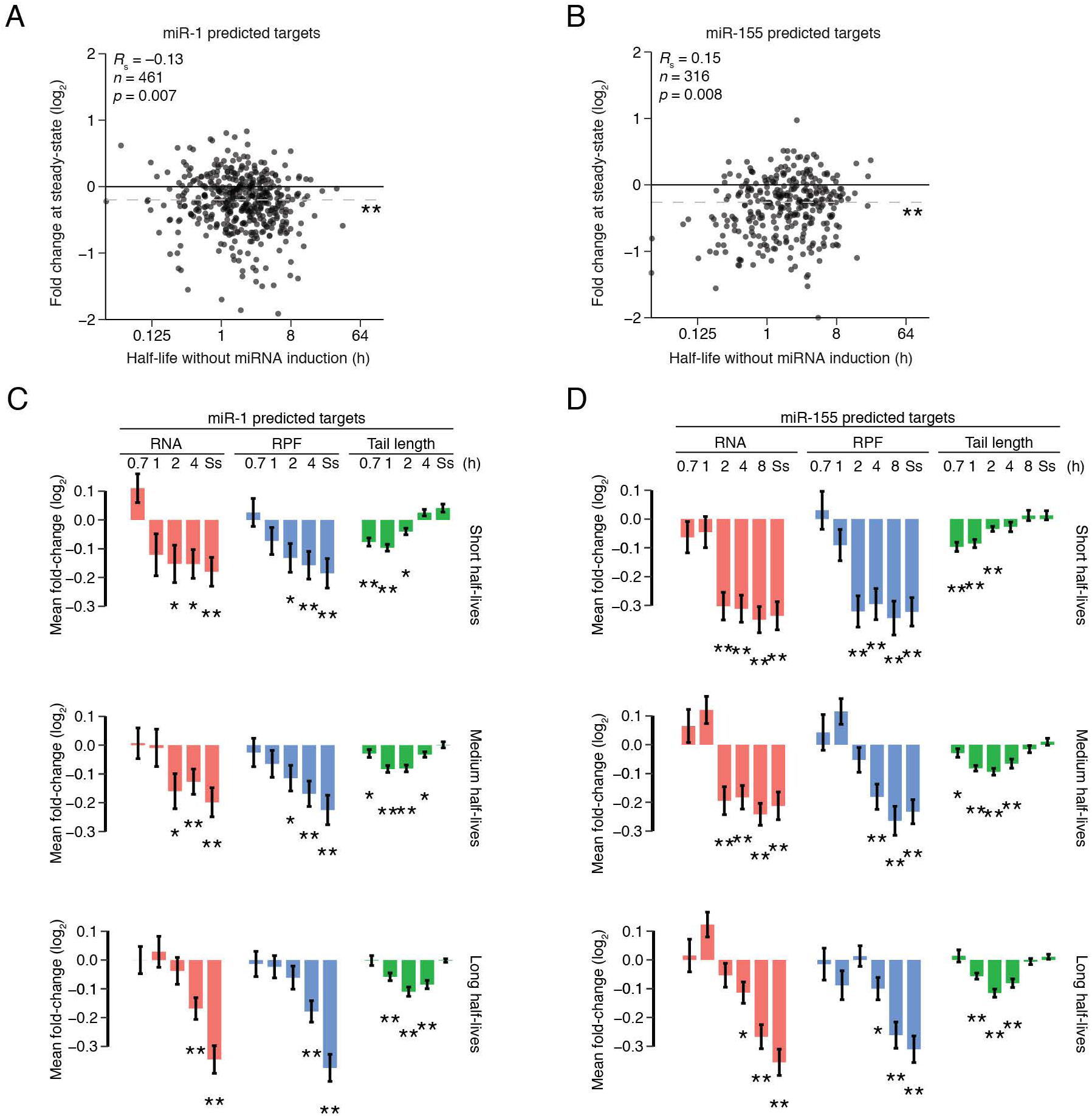
mRNA Half-life as a Determinant of miRNA-Mediated Repression. (A) The influence of basal half-lives on miR-1–mediated steady-state repression. miR-1– mediated changes in RNA abundance at steady state compared to mRNA half-life in the uninduced condition. The median fold-change of no-site mRNAs (n = 1274) was subtracted from the fold-changes of predicted targets (n = 461). Horizontal dashed grey lines denote the median values of each group, with asterisks denoting significance testing as in Figure 1A. The p value is the rank-order association test indicating the probability of a slope ≠ 0. (B) The influence of basal half-lives on miR-155–mediated steady-state repression. miR-155–mediated changes in RNA abundance at steady state compared to mRNA half-life in the uninduced condition, with no-site mRNA fold-changes (n = 1862) subtracted from the fold-change of the predicted targets (n = 316). Otherwise as in (A). (C) Dynamics of miR-1–mediated repression of predicted miR-1 targets split into three equally sized groups based on half-life. The groups have half-lives ranging from 0.01– 0.985 h, 0.985–2.365 h, and 2.365–20 h, respectively, with 92 target genes in each group, based on half-life measurements from uninduced miR-155–inducible cells. For each group, fold-changes were normalized to those of a set of no-site mRNAs with the same range of half-lives (n = 183, 168, and 289, respectively). Otherwise, as in (A). (D) Dynamics of miR-155–mediated repression of predicted miR-155 targets split into three equally sized groups based on half-life. These groups have half-lives ranging from 0.01–1.03 h, 1.03–2.8 h, or 2.8–20 h, respectively, with 83 target genes in each group, and were normalized to a set of no-site mRNAs with the same range of half-lives (n = 368, 462, and 614, respectively). Otherwise as in (A). See also Figure S4.

Our conclusion that the induced miRNAs acted together with basal decay factors to further reduce the stability of mRNAs that were already short lived predicted that miRNA effects would be observed earlier for short-lived mRNAs than for long-lived mRNAs. Analysis of our pre-steady-state results confirmed this prediction. Steady-state RNA and RPF repression levels were reached substantially earlier for predicted targets of miR-1 with shorter basal half-lives, with maximal miRNA-mediated tail shortening also occurring earlier for these predicted targets (Figure 5C). Although the dynamics of miRNA-mediated deadenylation and decay came into sharper focus when grouping predicted targets with similar basal half-lives, we still did not observe compelling evidence for miRNA-mediated translational repression, even at early time intervals. Analogous results were observed for the destabilization and tail-length dynamics of the top predicted targets (Figure S4A) and for targets and top predicted targets of miR-155 after inducing this miRNA (Figure 5D and S4B). Collectively, these analyses demonstrated that the basal half-life of the mRNA target strongly influences the dynamics of miRNA-mediated repression, again implying connections between the basal and miRNA decay factors.

## Discussion

Our experiments provided the opportunity to observe the early effects of miRNAs on translation and also learn how miRNAs influence the dynamics of deadenylation and decay in mammalian cells. Although in postembryonic cells miRNAs mostly act to enhance the decay of their mRNA targets (Guo et al., 2010; Eichhorn et al., 2014), translational repression can comprise a relatively greater fraction of the repression observed soon after a miRNA encounters an mRNA target. However, the magnitude of this phenomenon depends on the experimental approach. Substantial effects on translation are observed soon after inducing a reporter with sites to an expressed miRNA (Bethune et al., 2012; Djuranovic et al., 2012), whereas more modest effects on endogenous mRNAs are observed soon after inducing a miRNA (Eichhorn et al., 2014).

The results of our metabolic-labeling approach provided insight into the different effect sizes observed for the two previous experimental approaches. Like the miRNA-induction experiments, our metabolic-labeling experiments examined effects in 3T3 cells (indeed, in the same clonal lines as used in the previous miRNA-induction experiments) and monitored endogenous mRNAs rather than reporters. Like the reporter experiments, our metabolic-labeling experiments examined the early response to a preexisting, highly expressed miRNA, which presumably provides the best opportunity to observe the early effects of the miRNAs, which were already at full strength when the early measurements were taken. Despite this presumed greater sensitivity, our metabolic-labeling experiments were no better than the miRNA-induction experiments in detecting translational repression—indeed, they appeared to be somewhat worse. They did not detect convincing translational repression at any time interval (Figures 3 and 5), whereas the miRNA-induction experiments performed in the same cell lines seem to detect some translational repression after miR-155 induction (but not after miR-1 induction) (Eichhorn et al., 2014). This inability of our metabolic-labeling approach to detect translational repression at early time intervals indicated that the notion that translational repression constitutes a notable component of the effect soon after a miRNA encounters a target mRNA does not apply to miRNA-mediated repression of endogenous mRNAs—at least not in NIH 3T3 cells.

In contrast to translational repression, miRNA-mediated enhanced deadenylation was observed at early time intervals. Indeed, it was reliably detected prior to or coincident with accelerated mRNA degradation—results that integrated deadenylation and previously observed effects of miRNA-mediated repression. The influence on poly(A)-tails peaked at intermediate time points, and then disappeared at steady-state. Explaining this disappearance was our discovery that miRNAs not only accelerate deadenylation of their regulatory targets, but also accelerate the decay of these target mRNAs once their tails become shortened.

We speculate that accelerated decay of short-tailed mRNAs occurs through accelerated decapping. This accelerated decapping need not be through the direct contacts between TNRC6 (or AGO) and the decapping machinery or its coactivators. Instead, the changes that occur as the miRNA silencing complex helps remodel target mRNA–protein complexes to accelerate deadenylation might also recruit the decapping machinery or its coactivators. Indeed, physical connections between the Ccr4–Not deadenylase complex and the decapping complex (Haas et al., 2010; Ozgur et al., 2010; Jonas and Izaurralde, 2015) as well as the intracellular colocalization of these complexes (Parker and Sheth, 2007) could help link deadenylation with decapping.

Our concurrent analysis of the dynamics of cytoplasmic mRNA metabolism shows that mRNAs that undergo more rapid deadenylation generally undergo more rapid decay once their tails have become short (Eisen et al., 2019). This more rapid decay of short-tailed mRNAs that had previously been more rapidly deadenylated prevents a large buildup of short-tailed isoforms of rapidly deadenylated mRNAs, thereby enabling the broad range in mRNA deadenylation rates to impart a similarly broad range in mRNA decay rates (Eisen et al., 2019). These results observed when examining mRNA metabolism more generally—not just when looking at the effects of miRNAs—suggest that processes of deadenylation and short-tailed–mRNA decay are coupled for not only miRNA targets but also targets of other degradative pathways. Moreover, our finding that miRNAs work in conjunction with and reinforce these other degradative pathways reveals another dimension of interconnectivity that further supports this emerging view of broad integration of post-transcriptional regulatory processes. In this view, targeting by an added miRNA would accelerate the coherently coupled processes of deadenylation and short-tailed–mRNA decay already occurring for each of the cellular mRNAs.

## Acknowledgements

We thank J. Kwasnieski and other members of the Bartel lab for helpful discussions and the Whitehead Genome Technology Core for high-throughput sequencing. This research was supported by NIH grants GM061835 and GM118135 (D.P.B.) and an NSF Graduate Research Fellowship (T.J.E.). D.P.B. is an investigator of the Howard Hughes Medical Institute.

## Author Contributions

A.O.S., S.W.E., T.J.E., and D.P.B. conceived the project and designed the study. T.J.E., S.W.E., and A.O.S. performed the molecular experiments and analysis. T.J.E. performed the computational modeling. A.O.S., S.W.E., and T.J.E. drafted the manuscript, and T.J.E. and D.P.B. revised the manuscript with input from the other authors.

## Methods

### Cell culture

Clonal 3T3 cell lines engineered to express miR-1 or miR-155 upon doxycycline treatment were previously described (Eichhorn et al., 2014). Cells were grown at 37°C in 5% CO_2_ in DMEM supplemented with 10% BCS (Sigma-Aldrich) and 2 µg/mL puromycin. For metabolic-labeling time courses, cells from each line were plated onto 500 cm^2^ plates at 6.6 million cells per plate and cultured for two days such that they reached ∼70–80% confluency. Cells from each line were then re-plated in parallel for the uninduced and induced conditions, again at 6.6 million cells per 500 cm^2^ plate, with the uninduced cells cultured in DMEM with 10% BCS, and the induced cells cultured in DMEM with 10% BCS and 2 µg/mL doxycycline. After 48 h, growth media was supplemented with 5-ethynyluridine (5EU, Jena Biosciences) (Jao and Salic, 2008) at a final concentration of 400 µM, and cells were harvested after the desired labeling intervals (Figure 2A). Four plates were harvested for each 40 min time interval, three plates for each 1 h time interval, and two plates for each other time interval. A plate that had never received 5EU was harvested in parallel for each condition.

Cells were harvested at 4°C, washed twice with 50 mL ice-cold 9.5 mM PBS, pH 7.3 containing 100 µg/mL cycloheximide and then used to prepare cytoplasmically enriched lysate as described (Subtelny et al., 2014). An aliquot of cleared lysate was flash frozen for use in ribosome profiling, and the rest of the lysate was added to 5 volumes of TRI reagent (Ambion) and frozen at –80°C. Samples stored in TRI reagent were thawed at room temperature, and RNA was purified according to the manufacturer’s protocol and used for RNA-seq and PAL-seq. The non-5EU sample for the miR-1 cell line was labeled with 100 µM 4-thiouridine for 4 h prior to harvesting. 4-thiouridine has no known cross-reactivity with the reagents used to label 5EU.

### RNA standards

Two sets of tail-length standards (set 1 and set 3, Table S1) were described previously (standard mix 2 and standard mix 1) (Subtelny et al., 2014). The other set of standards (set 2, Table S1) was prepared based on a 705 nt fragment of the *Renilla* luciferase mRNA, which was transcribed and gel purified as described (Subtelny et al., 2014) and then capped using a Vaccinia capping system (2000 µL reaction containing 500 µg RNA, 1000 U Vaccinia capping enzyme (NEB), 1X Capping Buffer (NEB), 0.1 mM S-adenosyl methionine, 0.5 mM GTP, 50 nM [α–^32^P]-GTP, 2000 U SUPERaseIn (ThermoFisher) at 37°C for 1 h), monitoring the amount of incorporated radioactivity to ensure that capping was quantitative. Following the capping reaction, the 2′,3′ cyclic phosphate at the 3′ end was removed using T4 polynucleotide kinase (Subtelny et al., 2014). The capped, dephosphorylated product was joined by splinted ligation to each of seven different poly(A)-tailed barcode oligonucleotides (Subtelny et al., 2014). These seven 3′-ligation partners included 110 and 210 nt poly(A) oligonucleotides prepared as described (Subtelny et al., 2014), and five gel-purified synthetic oligonucleotides (IDT), one with a 10 nt poly(A) tract and the other four with a 29 nt poly(A) tract followed by either A, C, G, or U. Ligation products were gel purified, mixed in desired ratios, with final ratios of the different-sized species confirmed by analysis on a denaturing polyacrylamide gel.

Short and long standards were used to monitor enrichment of 5EU-containing fragmented RNA or non-fragmented RNA, respectively. Short 5EU standards were prepared by in vitro transcription of annealed DNA oligos to produce a 30 nt and 40 nt RNA, with the latter containing a single 5EU (Table S1). In vitro transcription was performed with the MEGAscript T7 transcription kit (ThermoFisher) according to the manufacturer’s protocol, except UTP was replaced with 5-ethynyluridine-triphosphate (Jena Biosciences) when transcribing the 40 nt RNA. Long standards were prepared by in vitro transcription of sequences encoding firefly luciferase and GFP using the MEGAscript T7 transcription kit and 0.1 µM PCR product as the template. When transcribing *GFP*, a 20:1 ratio of UTP to 5-ethynyluridine-triphosphate was used. Short and long standards were gel purified and stored at –80°C. Prior to use, a portion of each standard was cap-labeled and gel purified again, which enabled measurement of the recovery of the 5EU-containing standard relative to that of the uridine-only standard.

Three 28–30 nt RNAs (Table S1) were synthesized (IDT) for use as quantification standards in RNA-seq and ribosome-profiling libraries. These standards were gel purified, and 0.1 fmol of each was added to each sample immediately prior to library preparation.

### Biotinylation of 5EU labeled RNA

For RNA-seq libraries, poly(A) RNA was purified from 50 µg total RNA of the 40 min, 1, 2, and 4 h samples and 25 µg total RNA of the 8 h sample and non-5EU-labeled sample using oligo(dT) Dynabeads (ThermoFisher) according to manufacturer’s protocol. RNA was fragmented and 27–33 nt fragments were isolated as described (Subtelny et al., 2014), short standards that monitored 5EU enrichment were added, and then Cu(II) catalysis was used to biotinylate 5EU in a 20 µL reaction containing 50 mM HEPES, pH 7.5, 4 mM disulfide biotin azide (Click Chemistry Tools), 2.5 mM CuSO_4_, 2.5 mM Tris(3-hydroxypropyltriazolylmethyl)amine (THPTA, Sigma-Aldrich), and 10 mM sodium ascorbate, incubated at room temperature for 1 h. Reactions were stopped with 5 mM EDTA and then extracted with phenol-chloroform (pH 8.0). For the steady-state samples, 5 µg of RNA from the 40 min sample was poly(A) selected and fragmented, and size-selected 27–33 nt fragments were carried forward without enriching for 5EU.

For ribosome profiling libraries, 800 µL aliquots of lysate were digested with 0.7 U/µL RNAse I (Ambion) for 30 min at room temperature and then run on a sucrose gradient to purify monosomes (Subtelny et al., 2014). For each 40 min sample, monosomes from three aliquots were combined. For samples from all other time intervals, monosomes from two aliquots were combined, and for the non-5EU-labeled sample, only one aliquot was run. RPFs were released and purified (Subtelny et al., 2014), short standards used to monitor 5EU enrichment and recovery were added (using a 1:10 ratio of 5EU-containing standard to non-5EU-containing standard), and then click reactions were performed as above. For the steady-state samples, RPFs were isolated from one aliquot of the 40 min sample and carried forward without enriching for 5EU.

For PAL-seq, long standards used to monitor 5EU enrichment and recovery were added to total RNA (using a 1:10 ratio of 5EU-containing standard to non-5EU-containing standard), and samples were click labeled as above in reactions with 2.5 µg/µL RNA. For samples from the uninduced and induced miR-1 time courses, click reactions were performed with 800, 525, 350, 200, or 50 µg total RNA for the 40 min, 1 h, 2 h, 4 h, or non-5EU-labeled samples, respectively. For samples from the uninduced miR-155 time course, click reactions were performed with 500, 500, 250, 200, or 100 µg total RNA for the 40 min, 1 h, 2 h, 4 h, or 8 h samples, and for samples from the induced miR-155 time course, click reactions were performed with 400, 400, 250, 200, or 100 µg total RNA for the 40 min, 1 h, 2 h, 4 h, or 8 h samples, respectively. The non-fragmented miR-155 time course samples did not include a non-5EU-labeled sample (used solely to determine the background recovery in Figure S2).

### Purification of 5EU labeled RNA

For RNA-seq and ribosome-profiling samples, Dynabeads MyOne Streptavidin C1 beads (ThermoFisher) for each set of samples were combined and batch washed, starting with 200 µL of beads per reaction. Beads were washed twice with 1X B&W buffer (5 mM Tris-HCl, pH 7.5, 0.5 mM EDTA, 1 M NaCl and 0.005% Tween-20), twice with solution A (0.1 M NaOH, 50 mM NaCl), twice with solution B (0.1 M NaCl), and then twice with water, using for each wash a volume equal to that of the initial bead suspension. Following the last wash, beads were resuspended in an initial bead volume of 1X high salt wash buffer (HSWB, 10 mM Tris-HCl, pH 7.4, 1 mM EDTA, 0.1 M NaCl, 0.01% Tween-20) supplemented with 0.5 µg/mL yeast RNA (ThermoFisher) and incubated at room temperature for 30 min with end-over-end rotation, again using a volume equal to that of the initial bead suspension. Beads were then washed three times with 200 µL 1X HSWB per reaction and split for each reaction during the last wash. After the wash was removed, sample RNA resuspended in 200 µL 1X HSWB was added to blocked beads and incubated with end-over-end rotation at room temperature for 30 min. Beads were washed twice with 800 µL 50°C water, incubating at 50°C for 2 min for each wash, and then twice with 800 µL 10X HSWB. RNA was eluted from beads by incubating with 200 µL 0.5 M tris(2-carboxyethyl)phosphine (TCEP, Sigma-Aldrich) at 50°C for 20 min with end-over-end rotation. The initial eluate was collected, and beads were resuspended in 150 µL water and eluted again, combining the two eluates for each sample. RNA from the eluate was then ethanol precipitated using linear acrylamide as a carrier.

Purifications of non-fragmented RNA were performed as above, except bead volumes were adjusted based on estimates of the amount of labeled RNA in each sample. For the uninduced and induced miR-1 samples, 467, 452, 575, 598, and 250 µL streptavidin beads were used for the 40 min, 1 h, 2 h, 4 h, and non-5EU-labeled samples, respectively. For the uninduced miR-155 samples, 292, 431, 410, 598, and 500 µL of beads were used, and for the induced miR-155 samples, 234, 345, 410, 598, and 500 µL of beads were used for the 40 min, 1 h, 2 h, 4 h, and 8 h samples, respectively.

Pilot experiments designed to optimize the 5EU biotinylation and purification confirmed that RNAs containing at least one 5EU could be purified efficiently, with over 80% of a model RNA substrate containing a single 5EU becoming biotinylated in a 1 h reaction (Figure S2A). This high reaction efficiency was critical for both the ribosome-profiling and RNA-seq analyses, as RPFs and the RNA fragments from the paired RNA-seq libraries were only ∼30 nt long and estimated to typically contain at most a single 5EU. Indeed, for each of the three protocols, which started with either full-length RNA (PAL-seq) or fragmented RNA (RNA-seq and ribosome-footprint profiling), captured labeled RNA was substantially enriched above background (Figure S2B and C).

### PAL-seq

We used an improved form of PAL-seq called PAL-seq v2, using RNA standards of defined tail lengths to monitor library preparation, sequencing, and the computational pipeline. Steady-state RNA (25 µg of unselected RNA from the 40 min sample) or half of the RNA eluted from each 5EU-selected sample was used to prepare PAL-seq libraries. Tail-length standard mixes (1 ng of set 1 and 2 ng of set 2 for each 5EU-selected sample, and twice these amounts for the steady-state sample), and trace 5′-radoiolabeled marker RNAs (Table S1) were added to each sample to assess tail-length measurements and ligation outcomes, respectively. Polyadenylated ends including those with a terminal uridine were ligated to a 3′-biotinylated adapter DNA oligonucleotide (1.8 µM) in the presence of two splint DNA oligonucleotides (1.25 µM and 0.25 µM for the U and A-containing splint oligos, respectively, Table S1) using T4 Rnl2 (NEB) in an overnight reaction at 18°C. Following 3′-adapter ligation the RNA was extracted with phenol-chloroform (pH 8.0), precipitated, resuspended in 1X RNA T1 sequence buffer (ThemoFisher), heated to 50°C for 5 min and then put on ice. RNase T1 was then added to a final concentration of 0.006 U/µL and the reaction incubated at room temperature for 30 min followed by phenol chloroform extraction and precipitation. RNA was subsequently captured on streptavidin beads, 5′ phosphorylated, and ligated to a 5′ adapter as described (Subtelny et al., 2014) but using a modified 5′ adapter sequence (Table S1). Following reverse transcription using SuperScript III (Invitrogen) with a barcode-containing DNA primer, cDNA was purified as described (Subtelny et al., 2014), except a 160–810 nt size range was selected. Libraries were amplified by PCR for 8 cycles using Titanium Taq polymerase according to the manufacturer’s protocol with a 1.5 min combined annealing/extension step at 57°C. PCR-amplified libraries were purified using AMPure beads (Agencourt, 40 µL beads per 50 µL PCR, two rounds of purification) according to the manufacturer’s instructions.

The use of a splinted ligation of the 3′ adapter to the poly(A) tail had the advantage of specifically ligating to mRNAs without the need to deplete ribosomes or other abundant RNAs. However, this approach was not suitable for acquiring measurements for mRNAs with tails that were either very short (< 8 nt) or extended by more than one uridine, because such tails would ligate less efficiently (or not at all) when using a splinted ligation to the 3′ adapter. To account for these mRNAs with either very short or highly modified tails, we implemented a protocol that used single-stranded (ss) ligation and different mRNA enrichment steps to prepare libraries from steady-state RNA isolated from miR-1 and miR-155 cell lines in uninduced and induced conditions. For each sample, 5 µg of total RNA was depleted of rRNA using RiboZero Gold HMR (Illumina) and further depleted of the 5.8s rRNA by subtractive hybridization. Subtractive hybridization was performed by mixing 2x SSC buffer (3M sodium chloride, 300mM sodium citrate, pH 7.0), total RNA, and 4.8µM of each 5.8s subtractive-hybridization oligo (Table S1) in a 50 µL reaction, heating the reaction to 70°C for 5 min, then cooling it at 1°C/min to 37°C to anneal the oligos to the RNA. During this cooling, 250 µL of Dynabeads MyOne Streptavidin C1 beads per sample (ThermoFisher) were washed twice with 1X B&W buffer (5 mM Tris-HCl, pH 7.5, 0.5 mM EDTA, 1 M NaCl and 0.005% Tween-20), twice with solution A (0.1 M NaOH, 50 mM NaCl), twice with solution B (0.1 M NaCl), and then resuspended in 50 µL of 2X B&W buffer. After cooling, the entire 50µL RNA/oligo mixture was added to 50 µL of washed beads, then incubated at room temperature for 15 min with end-over-end rotation. The sample was then magnetized and the supernatant was withdrawn and precipitated by adding 284 µL of water, 4 µL of 5 mg/mL linear acrylamide, and 1 mL of ice-cold 96% ethanol. After resuspension, RNA was ligated to a 3′ adapter containing four random-sequence nucleotides and an adenylyl group at its 5′ end (Table S1) in a 70 µl reaction containing 10 µM adapter, 1X T4 RNA Ligase Reaction Buffer (NEB), 20 U/µL T4 RNA Ligase 2 truncated KQ (NEB), 0.3 U/µL SUPERaseIn (ThermoFisher), and 20% PEG 8000. The reaction was incubated at 22°C overnight and then stopped by addition of EDTA (3.5 mM final after bringing the reaction to 400 µL with water). RNA was phenol–chloroform extracted, precipitated, and subsequent library preparation was as for the splinted-ligation libraries.

PAL-seq v2 libraries were sequenced on an Illumina HiSeq 2500 operating in rapid mode. Hybridization mixes were prepared with 0.375 fmol PCR-amplified library that had been denatured with standard NaOH treatment and brought to a final volume of 125 µL with HT1 hybridization buffer (Illumina, 3 pM library in final mix). Following standard cluster generation and sequencing-primer hybridization, two dark cycles were performed for the splint-ligation libraries (i.e., two rounds of standard sequencing-by-synthesis in which imaging was skipped), which extended the sequencing primer by 2 nt, thereby enabling measurement of poly(A) tails terminating in non-adenosine bases. For the direct-ligation libraries, six dark cycles were performed instead of two, which extended the sequencing primer past the four random-sequence nucleotides in the 3′ adapter and then the last two residues of the tail.

Following the two dark cycles, a custom primer-extension reaction was performed on the sequencer using 50 µM dTTP as the only nucleoside triphosphate in the reaction. To perform this extension, the flow cell temperature was first set to 20°C. Then, 120 µL of universal sequencing buffer (USB, Illumina) was flowed over each lane, followed by 150 µL of Klenow buffer (NEB buffer 2 supplemented with 0.02% Tween-20). Reaction mix (Klenow buffer, 50 µM dTTP, and 0.1 U/µL units Large Klenow Fragment, NEB) was then flowed on in two aliquots (150 µL and 100 µL). The flow-cell temperature was then increased to 37°C at a rate of 8.5°C per min and the incubation continued another 2 min after reaching 37°C. 150 µL of fresh reaction mix was then flowed in, and following a 2 min incubation, 75 µL of reaction mix was flowed in eight times, with each flow followed by a 2 min incubation. The reaction was stopped by decreasing the flow cell temperature to 20 °C, flowing in 150 µL of quench buffer (Illumina HT2 buffer supplemented with 10 mM EDTA) and then washing with 75 µL of HT2 buffer. The flow cell was prepared for subsequent sequencing with a 150 µL and a 75 µL flow of HT1 buffer (Illumina). 50 cycles of standard sequencing-by-synthesis were then performed to yield the first sequencing read (read 1). XML files used for this protocol are provided at https://github.com/kslin/tail-seq.

The flow cell was stripped, a barcode sequencing primer was annealed, and seven cycles of standard sequencing-by-synthesis were performed to read the barcode. The flow cell was then stripped again, and the same primer as used for read 1 was hybridized and used to prime 250 cycles of standard sequencing-by-synthesis to generate read 2. Thus, each PAL-seq tag consisted of three reads: read 1, read 2, and the indexing (barcode) read. For cases in which a tag corresponded to a polyadenylated mRNA, read 1 was the reverse complement of the 3′ end of the mRNA immediately 5′ of the poly(A) tail and was used to identify the mRNA and cleavage-and-polyadenylation site of long-tailed mRNAs. The indexing read was used to identify the sample, and read 2 was used to measure poly(A)-tail length and identify the mRNA and cleavage-and-polyadenylation site of short-tailed mRNAs. The intensity files of reads 1 and 2 were used for poly(A)-tail length determination, along with the Illumina fastq files.

### PAL-seq v2 data analysis

Tail lengths for the splinted-ligation data were determined using a Gaussian hidden Markov model (GHMM) from the python2.7 package ghmm (http://ghmm.org/), analogous to the model used in TAIL-seq (Chang et al., 2014) and described in the next paragraph. Read 1 was mapped using STAR (v2.5.4b) run with the parameters ‘-- alignIntronMax 1 --outFilterMultimapNmax 1 --outFilterMismatchNoverLmax 0.04 -- outFilterIntronMotifs RemoveNoncanonicalUnannotated --outSJfilterReads’, aligning to an index of the mouse genome built using mm10 transcript annotations that had been compressed to unique instances of each gene selecting the longest transcript and removing all overlapping transcripts on the same strand (Eichhorn et al., 2014). The genome index also included sequences of the quantification spikes and the common portion of the poly(A)-tail length standards. The sequences that identified each RNA standard (the last 20 nt of each standard sequence, Table S1) were not aligned using STAR. Instead, the unix program grep (v2.16) was used to determine which reads matched each standard (allowing no mismatches), and these reads were added to the aligned reads from the STAR output. Tags corresponding to annotated 3′ UTRs of mRNAs were identified using bedtools (v2.26.0), and if the poly(A)-tail read (read 2) contained a stretch of ≥ 10 T residues (the reverse complement of the tail) in an 11-nt window within the first 30 nt, this read was carried forward for GHMM analysis. If read 2 failed to satisfy this criterion but began with ≥ 4 T residues, the tail length was called based on the number of contiguous T residues at the start of read 2; by definition, these tails were < 10 nt and thus easily determined by direct sequencing.

For each read 2 that was to be input into the GHMM a ‘T signal’ was first calculated by normalizing the intensity of each channel for each cycle to the average intensity of that channel when reading that base in read 1 and then dividing the thymidine channel by the sum of the other three channels. Sometimes a position in a read would have a value of 0 for all four channels. A read was discarded if it contained more than five such positions. Otherwise, the values for these positions were imputed using the mean of the five non-zero signal values upstream and downstream (ten positions total) of the zero-valued position. A three-state GHMM was then used to decode the sequence of states that occurred in read 2. It consisted of an initiation state (state 1), a poly(A)-tail state (state 2), and a non-poly(A)-tail state (state 3). All reads start in state 1. From state 1 the model can remain in state 1 or transition to state 2. From state 2 the model can either remain in state 2 or transition to state 3. The model was initialized with the following transition probabilities:

| *from \ to* | *state<sub>1</sub>* | *state<sub>2</sub>* | *state<sub>3</sub>* |
| --- | --- | --- | --- |
| *state<sub>1</sub>* | 0.001 | 0.95 | 0.049 |
| *state<sub>2</sub>* | 0.001 | 0.95 | 0.049 |
| *state<sub>3</sub>* | 0.001 | 0.001 | 0.998 |

The initial emissions were Gaussian distributions with means of 100, 1, and –1 and variances of 1, 0.25 and 0.25, respectively. In general, the emission Gaussians for the model corresponded to the logarithm of the calculated T signal at each sequenced base in read 2. The initial state probabilities were 0.998, 0.001, and 0.001 for states 1, 2 and 3, respectively.

After initializing the model, unsupervised training was performed on 10,000 randomly selected PAL-seq tags, and then the trained model was used to decode all tags, with the number of state 2 cycles reporting the poly(A)-tail length for a tag. Only genes with ≥ 50 poly(A)-tail length measurements were considered for analyses involving mean poly(A)-tail lengths.

### Analysis of PAL-seq data from the ss-ligation protocol

To account for mRNAs with very short tails or extensive terminal modifications, we implemented a version of PAL-seq that did not use splinted ligation. Tail lengths from these ss-ligation datasets, acquired for steady-state samples from both cell lines, were determined using a modified version of the PAL-seq analysis pipeline written for python3. The T-signal in this pipeline was modified to allow more accurate quantification of 0-length tails. Instead of normalizing the intensity of each channel for each cycle to the average intensity of that channel when reading that base in read 1, the intensity of each channel was normalized to the average intensity of the channels for the other three bases in read 1. The intensity of the T channel was then divided by the sum of the other channel intensities to calculate the T signal, and tails were called using the hmmlearn package (v0.2.0). Tags representing short tails, including short tails that ended with many non-A residues, were identified as those for which read 1 and read 2 mapped to the same mRNA 3′ UTR (usually ∼4% of the tags). Tail lengths for these tags were called without the use of the GHMM. Instead, their tail lengths were determined by string matching, allowing any number of untemplated U residues but no more than two G or C residues to precede the A stretch. Tags not identified as representing short-tails were analyzed using the GHMM, excluding from further analysis occasional outliers determined by the GHMM to have tails ≤ 8 nt.

Most of the tags that had either only a very short tail or no tail did not correspond to mRNA cleavage-and-polyadenylation sites. Therefore, to be carried forward in our analysis, short-tailed tags were required to have a 3′-most genome mapping position (as determined from read 1 but requiring that read 2 also map uniquely to the same 3′ UTR) that fell within a 10 nt window of a PAL-seq–annotated cleavage-and-polyadenylation site.

Although the single-stranded ligation protocol provided the opportunity to account for mRNAs with very short or highly modified tails, examination of the recovery of internal standards indicated that tags representing longer tails (≥ 100 nt) were not as well recovered in the datasets in which we implemented ss ligation. Therefore, for steady-state samples from each cell line, we generated composite tail-length distributions in which the ss-ligation dataset contributed to the distribution of tails < 50 nt, and the splint-ligation dataset contributed to the distribution of tails ≥ 50 nt. For example, *Slc38a2* had 635 standard PAL-seq tags, 169 of which (∼27%) had tails < 50 nt, and this same gene had 703 ss-ligation PAL-seq tags, 393 of which (∼56%) had tails < 50 nt. The composite tail-length distribution replaced the 169 short-tailed splint-ligation PAL-seq tags with the 393 short-tailed ss-ligation PAL-seq tags, normalizing the latter cohort by a scaling factor. This scaling factor was determined from the ratio of the counts of the splint-ligation tags with tail lengths between 30–70 nt (135 tags) to the counts of the corresponding tags in the ss-ligation dataset (153 tags).

3′-end annotations were generated from PAL-seq tags with tails ≥ 11 nt, using an algorithm previously developed for data from poly(A)-position profiling by sequencing (3P-seq) (Jan et al., 2011). Each PAL-seq read 1 that mapped (with at least 1 nt of overlap) to an annotated 3′ UTR (Eichhorn et al., 2014) was compiled by the genomic coordinate of its 3′-most nucleotide. The position with the most mapped reads was annotated as a 3′ end. All reads within 10 nt of this end (a 21 nt window) were assigned to this end and removed from subsequent consideration. This process was repeated until there were no remaining 3′ UTR-mapped reads. For each gene, the 3′-end annotations were used in subsequent analyses if they accounted for ≥ 10% of the 3′ UTR-mapping reads for that gene.

Documentation and code to calculate and analyze T signals and determine tail lengths are available for both the splint-ligation and ss-ligation pipelines at https://github.com/kslin/PAL-seq.

### RNA-seq and ribosome profiling

Fragmented poly(A)-selected RNA and RPFs were supplemented with three short quantification standards (Table S1), and then ligated to adaptors, reverse-transcribed, and amplified to prepare the RNA-seq and ribosome-profiling libraries, respectively (Subtelny et al., 2014). These libraries were sequenced on an Illumina HiSeq 2500. For all RNA-seq and ribosome-profiling data, only reads mapping to ORFs of annotated gene models (Eichhorn et al., 2014) were considered, excluding the first 50 nt of each ORF. A cutoff of ≥ 10 reads per million mapped reads (RPM) was applied to each sample, with the exception of those samples generated from miRNA-induced cell lines, for which no cutoff was applied.

### Calculation of miRNA-mediated repression

Secondary effects of expressing a miRNA can have a greater impact on mRNAs with longer 3′ UTRs relative to those with shorter 3′ UTRs (Agarwal et al., 2015), presumably because longer 3′ UTRs tend to contain more sites to other regulatory factors, including other miRNAs. As a result, 3′-UTR length differences can complicate the measurement of the repressive effects of an expressed miRNA. For this reason, we first normalized the fold-changes of all mRNAs based on their 3′ UTR length. The relationship between the fold-change for all mRNAs without a 6-nt seed-matched site to the induced miRNA in the entire transcript (no-site mRNAs) and 3′ UTR length was calculated using linear regression, and then the fold-changes of all mRNAs (with and without a target site) were normalized by their 3′ UTR lengths such that the slope of the relationship between no-site mRNAs and 3′ UTR length was 0. We then compared normalized fold-changes for mRNAs containing at least one predicted miRNA target site in their 3′-UTR to those for the no-site mRNAs. For all mRNAs passing our expression threshold in the uninduced samples, we calculated the log_2_ fold-changes in mRNA abundance, RPF abundance, or poly(A)-tail length in samples from induced cells relative to the corresponding samples from uninduced cells. The repressive effect of the miRNA on a set of predicted miRNA targets was then calculated by subtracting the median-normalized fold-change for no-site mRNAs from the mean-normalized fold-change for a set of predicted targets. Top targets were defined using RPF measurements from a previous study (Eichhorn et al., 2014), choosing from among the predicted targets those with expression that decreased to ≤ 75% of their original expression after 12 hours of miRNA induction.

The cumulative-distribution plots each used a subset of all possible no-site mRNAs for display and analysis. The subsets were generated by choosing, for each site-containing mRNA, a no-site mRNA from the complete list of no-site mRNAs by random sampling (without replacement) such that the chosen no-site mRNA had a UTR length that fell within 20% of the length of the UTR of the site-containing mRNA. For each dataset, this sampling was performed 101 times and the control cohort yielding the median fold-change was chosen and used for plotting and analysis. The residual pool of no-site mRNAs was occasionally not sufficient to choose a no-site mRNA for each site-containing mRNA, which is why, for some figures, the no-site cohort had a slightly smaller n, particularly when the site-containing cohort had a large n.

### Modeling miRNA effects

To fit miRNA effects, we used model parameters for mRNAs from each gene, which had been fit for the uninduced miR-1 and miR-155 cell lines (Eisen et al., 2019), as a starting point for an additional round of model fitting. In this fitting, either the deadenylation rate constant, the decapping rate constant, or both of these rate constants were tuned in order to minimize the residual between the observed and simulated miRNA-induced fold-changes in mRNA abundance at steady state (calculated by summing the output of the model over all tail lengths ≤ 249). The optimization was performed using the L-BFGS-B method of the optim function in R.

### Background subtraction and normalization for PAL-seq data

Although the efficacy of the 5EU purification enabled efficient enrichment of labeled RNAs at short time intervals, we also modeled and corrected for residual background caused by non-specific binding of the unlabeled RNA to the streptavidin beads (Eisen et al., 2019).

We designed our background model under the assumption that the background in the time courses stems primarily from the capture of a fixed amount of non-5EU labeled mRNA during the 5EU purification. Accordingly, we subtracted a fraction (0.3%) of the steady-state data from each continuous-labeling dataset. This fraction of input sample was chosen such that at 40 min long-lived genes (half-life ≥ 8 h) had no mRNAs with tail lengths ≤ ∼100 nt on average, but short-lived genes (half-life ≤ 30 min) were unaffected (Eisen et al., 2019). The fraction of each input sample to subtract was chosen such that at 0 h long-lived genes (half-life ≥ 8 h) had no mRNAs with tail lengths ≤ ∼100 nt on average, but short-lived genes (half-life ≤ 30 min) were unaffected. Genes were included in the final background-subtracted set only if the sum of their background-subtracted tag counts was ≥ 50 tags.

After background subtraction, PAL-seq datasets were scaled to each time interval by matching the total number of background-subtracted tags for all genes at all tail lengths to the total number of tags for all genes for the corresponding time interval in the RNA-seq data. The scaled PAL-seq data were then used to compute half-lives for each gene, scaling the steady-state sample using a globally fitted constant.

### Accession numbers

Raw and processed RNA-seq, ribosome profiling, and PAL-seq data is available at the GEO, accession number GSE134660. Code for configuring an Illumina HiSeq 2500 machine for PAL-seq and for calculation of tail lengths from PAL-seq data is available at https://github.com/kslin/PAL-seq.

## Supplemental Figure Legends

**Figure S1.**
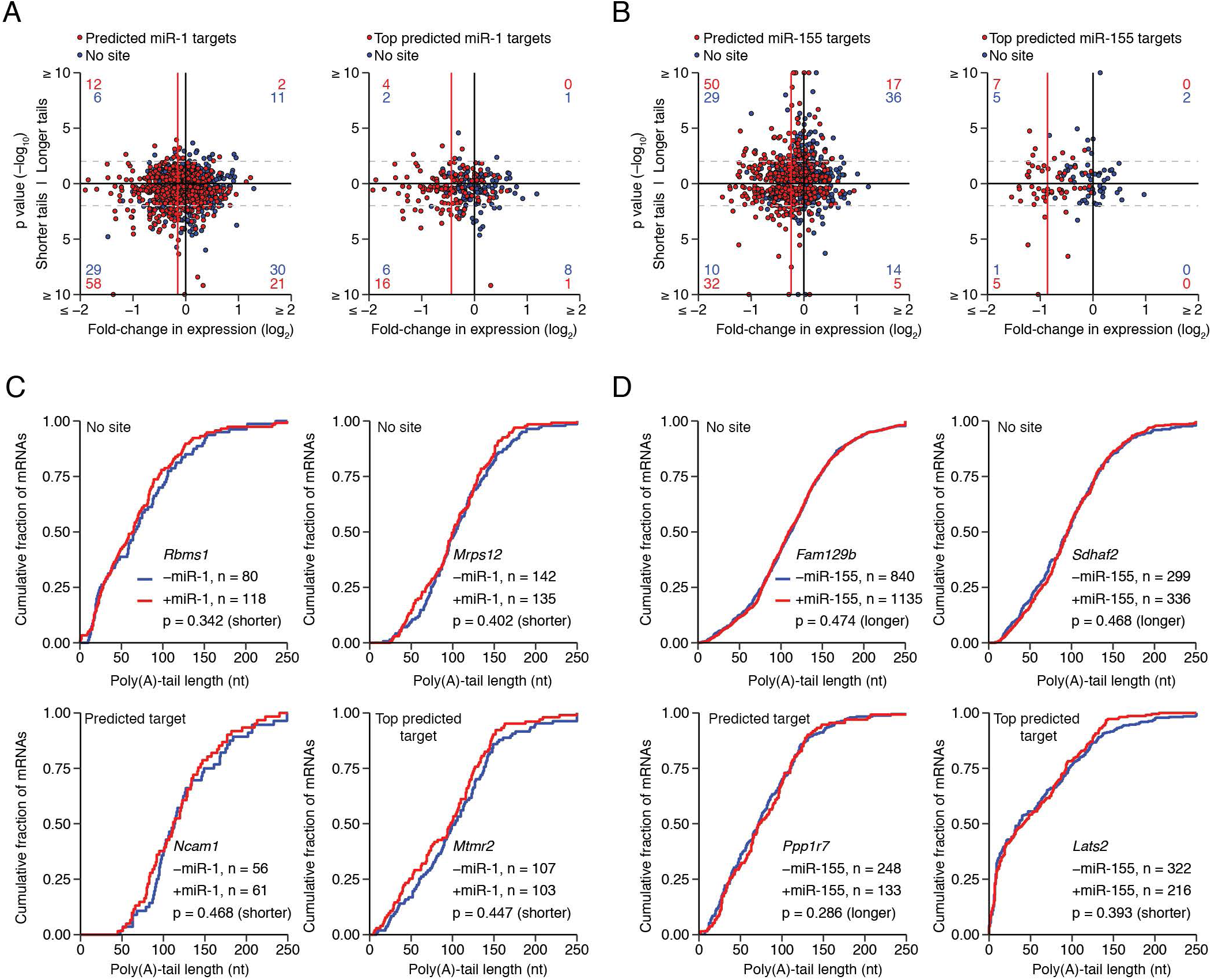
Very Small Effect of miRNAs on Steady-State Poly(A)-Tail Lengths When Examining mRNAs from Individual Genes, Related to Figure 1. (A) Influence of miR-1 on tail-length distributions of mRNAs from individual genes. On the left, K–S tests were performed for each predicted target of miR-1 (n = 643, red) and a corresponding set of no-site mRNAs (n = 624, blue). The p values (–log_10_) from these tests are plotted as a function of the miR-1–mediated fold-change in expression (log_2_), with placement above or below the *x* axis indicating the direction of the more significant p value in a one-tailed test, with values above the axis indicating a more significant increase in tail lengths with induction of the miR-1 and values below the axis indicating a more significant decrease in tail lengths. Horizontal dashed lines indicate p values of 0.01; for each quadrant, the number of predicted-target and no-site genes with p values below this threshold is indicated (red and blue, respectively). The median of the fold-change in expression for predicted targets of miR-1 after subtracting the median fold-change for the no-site controls is also indicated (red line). The analysis on the right is the same but considering the top predicted targets and their no-site controls (n = 130 and 130, respectively). (B) Influence of miR-155 on tail-length distributions of mRNAs from individual genes. Analysis is as in (A), except for predicted targets of miR-155 and their no-site controls (n = 417 and 417, respectively) and top predicted targets of miR-155 and their no-site controls (n = 57 and 57, respectively). (C) Typical influence of miR-1 on tail-length distributions. Tail-length distributions observed with and without miRNA induction (red and blue, respectively) are plotted for mRNAs with the median p values in (A). *Rbms* (top left) was the mRNA from the no-site cohort of the predicted miR-1 targets with the median p value; *Mrps12* (top right) was the mRNA from the no-site cohort of the top predicted miR-1 targets with the median p value; *Ncam1* was the predicted miR-1 target with the median p value, and *Mtmr2* was the top predicted miR-1 target with the median p value. The p values, calculated in (A), are shown along with the numbers of tags in each distribution (key). (D) Typical influence of miR-155 on tail-length distributions. Tail-length distributions observed with and without miRNA induction (red and blue, respectively) are plotted for mRNAs with the median p values in (B). Otherwise, this panel is as in (A).

**Figure S2.**
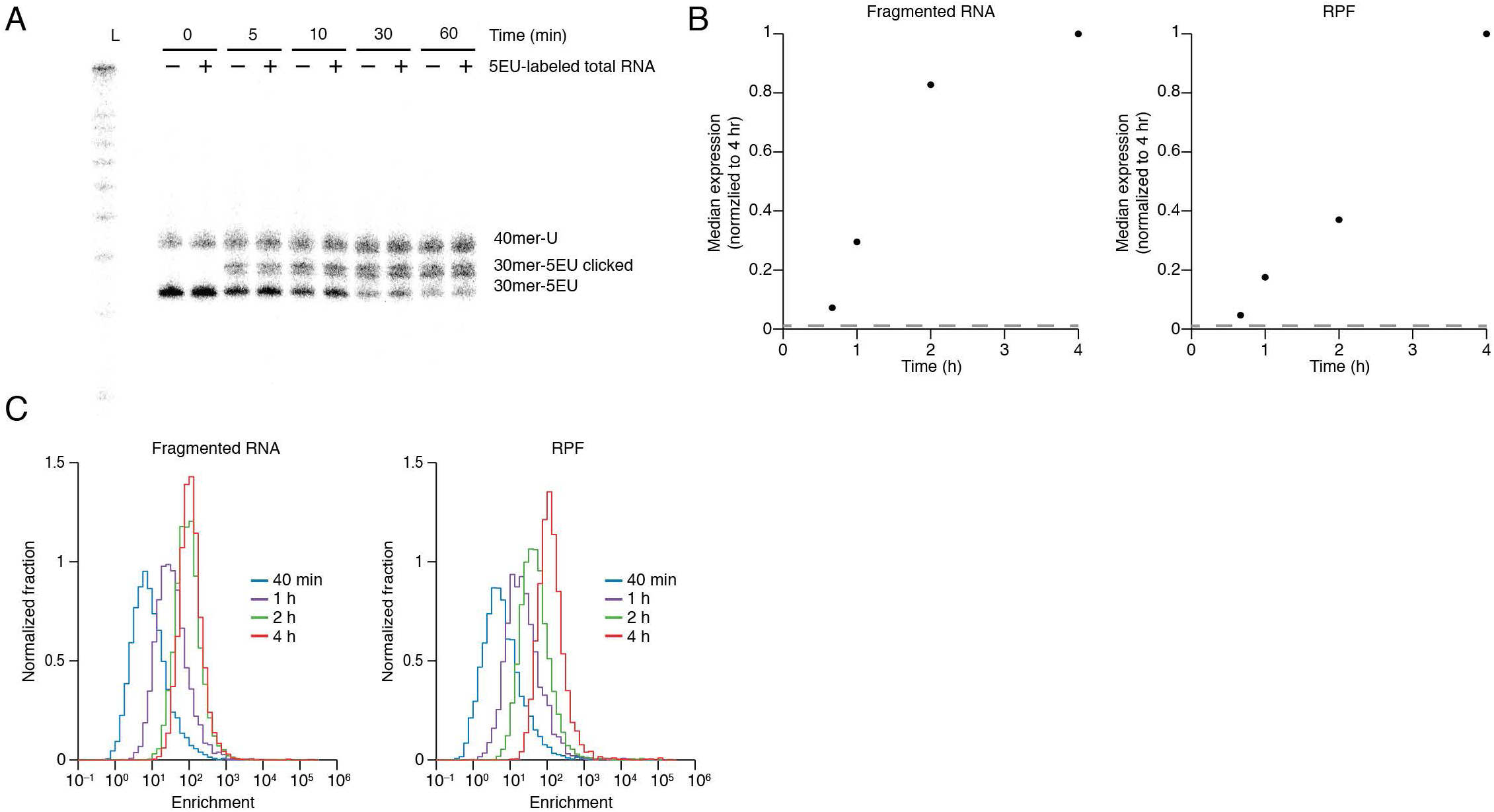
Enrichment of 5EU–labeled RNA, Related to Figure 2. (A) Biotinylation of a 5EU–containing RNA. A 40 nt RNA containing a single uridine and a 30 nt RNA containing a single 5EU were mixed with or without an additional 50 µg of 5EU–labeled total RNA and then reacted with disulfide biotin azide for the indicated amount of time. Decreased mobility was observed upon biotinylation, with ∼80% of the 5EU–containing RNA shifting to a slower mobility (30mer-5EU clicked) after 1 h and no detectable formation of a slower-mobility product from the 40 nt RNA that lacked 5EU. The clicked-labeled form of the RNA tended to run as a doublet, but reduced to a single band upon treatment with TCEP (not shown). Note that although in this pilot experiment the 40 nt RNA lacks 5EU and the 30 nt RNA contains it, the samples prepared for sequencing used a 30 nt RNA lacking 5EU and a 40 nt RNA containing it so that the 5EU containing RNA would be removed after size selection. (B) Abundance normalization of sequencing reads. mRNA was fragmented using either alkali (left) or as ribosome protected fragments (RPFs, right), and then 5EU-containing fragments were isolated and used to make libraries. To evaluate the background in labeled samples, we also attempted to isolate 5EU-containing fragments of both types starting with RNA from cells that had never received 5EU. All samples contained RNA standards that were used to scale mRNA abundance measurements. Median-scaled expression is plotted for both sets of libraries in the miR-1 line after normalizing to that of the 4 h sample for mRNAs passing cutoffs for RNA-seq and RPF analyses. No cutoff and a pseudo-count of 0.1 reads was applied to the unlabeled samples used to assess background. The normalized background expression is plotted as a dashed grey line. (C) Enrichment of labeled RNAs in fragmented RNA and RPF samples. Absolute mRNA abundance in each sample was normalized to expression in the corresponding unlabeled background sample, and the enrichment ratios for mRNA from each gene are plotted as a histogram. An enrichment of 1 would indicate as many reads in the labeled sample as in the background sample. For all mRNAs with enrichments > 1, most reads came from labeled RNA; otherwise, as in (B).

**Figure S3.**
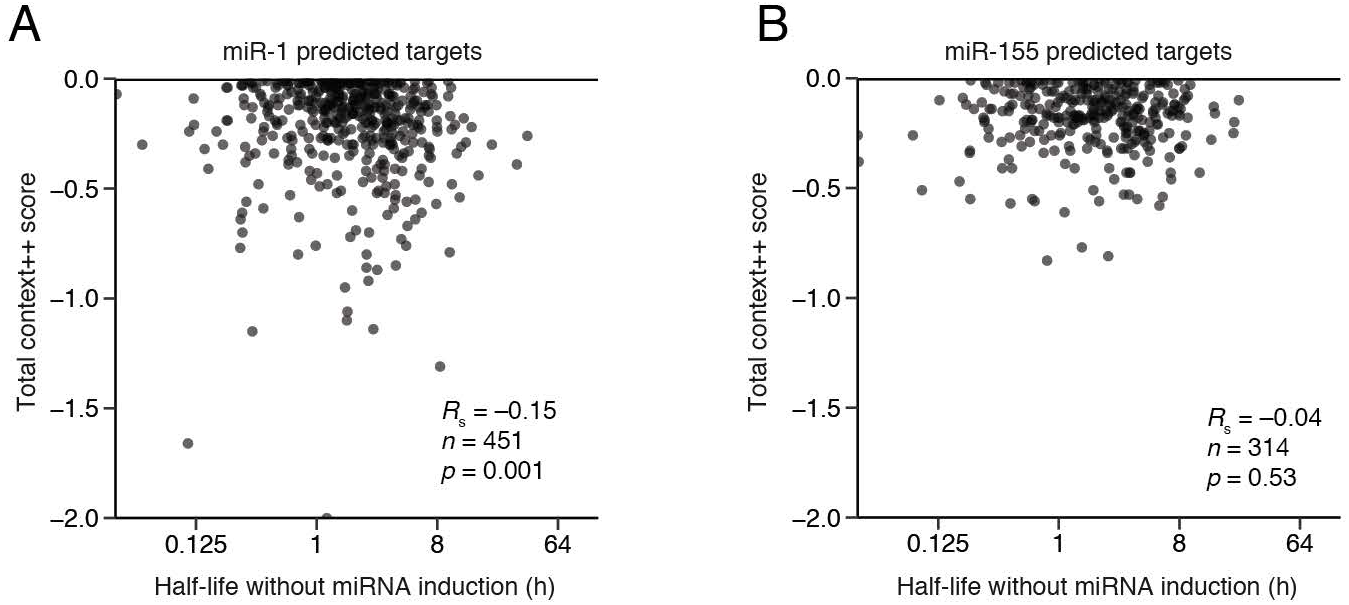
Half-life Analyses of miRNA Top Targets, Related to Figure 5. (A) The relationship between basal half-lives and predicted miR-1 target-site efficacy. Total context++ scores (Agarwal et al., 2015) for predicted targets of miR-1 were plotted with respect to half-life in the uninduced condition. mRNAs with predicted sites that do not have context ++ scores (n = 10) have sites in mRNA isoforms that fell below the TargetScan7 cutoff for isoform abundance. Otherwise as in Figure 5A. (B) The relationship between basal half-lives and predicted miR-155 target-site efficacy. As in (A), except plotting the miR-155 predicted targets in Figure 5B and the respective context++ scores. Two mRNAs had sites in isoforms that fell below the TargetScan7 cutoff for isoform abundance.

**Figure S4.**
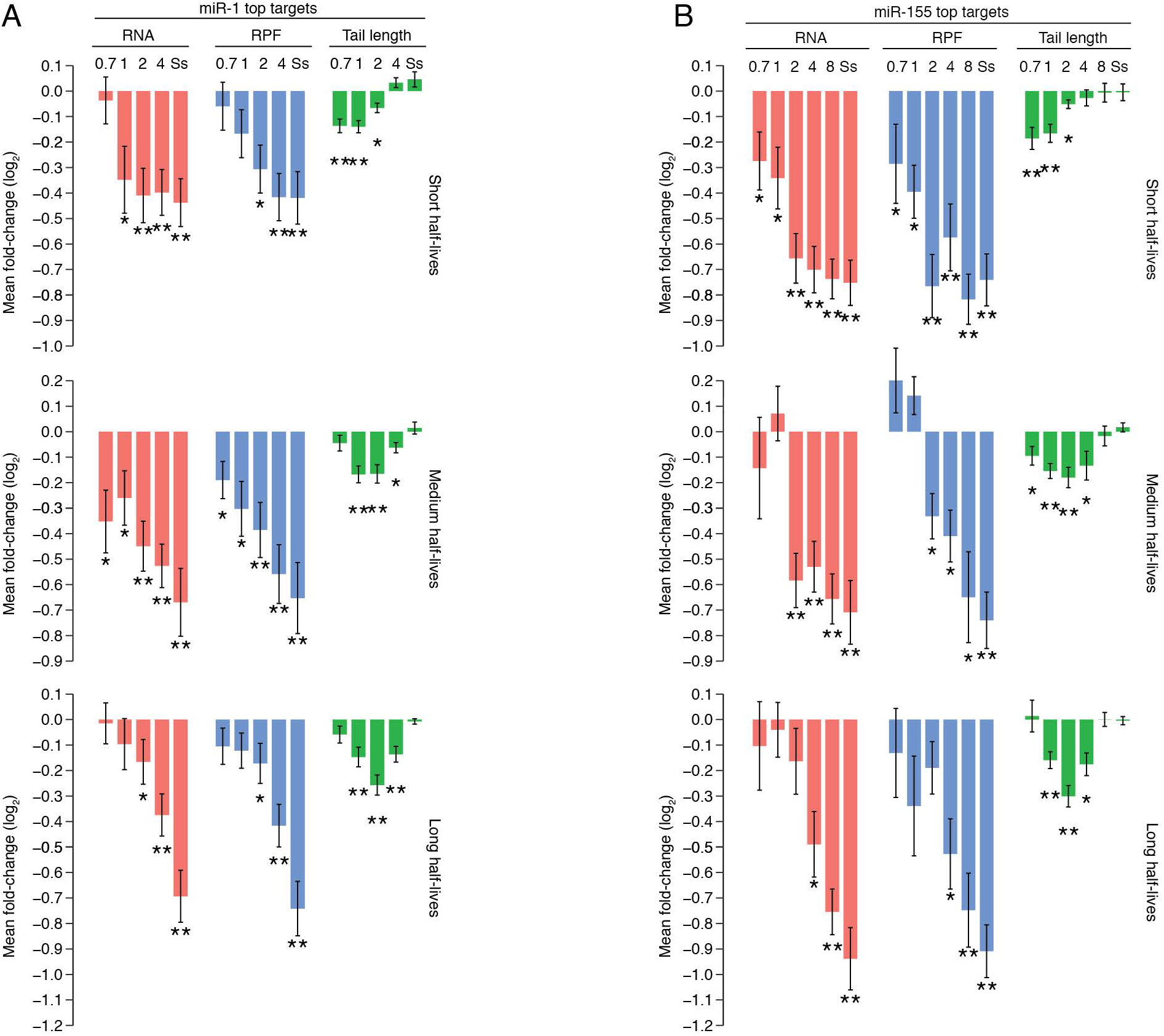
Half-life Analyses of miRNA Top Targets, Related to Figure 5. (A) The dynamics of miR-1–mediated repression for top targets. miR-1-mediated changes in RNA and RPF abundance and mean poly(A)-tail length for top miR-1 targets that were split into three groups (n = 25, 17 and 24 for short, medium and long half-life targets, respectively, using the same no-site genes and half-life ranges as in Figure 4B). Only mRNAs passing cutoffs for half-life measurements and RNA, RPF, and poly(A)-tail length measurements at all time points were analyzed. (B) The dynamics of miR-155–mediated repression for top targets. miR-155–mediated changes in RNA and RPF abundance and mean poly(A)-tail length for top miR-155 targets that were split into three groups (n = 19, 11 and 11 for short, medium and long half-life targets, respectively). Half-life cutoffs and numbers of no-site genes as in Figure 4C. Otherwise, as in (A).

